# User-friendly scheduler Using a hybrid architecture and supercomputing for big data processing

**DOI:** 10.1101/2025.09.01.673517

**Authors:** Patrick McKeever, Varun Mittal, Bryce Fukuda, Ka Yee Yeung, Ling-Hong Hung

## Abstract

The exponential growth of omics data requires novel strategies for storage, transfer, and processing of said data. We present a scheduler based on the Temporal.io workflow framework which enables two key optimizations of bioinformatics workflows. Firstly, we enable users to transparently map workflow steps to diverse execution environments, including high-performance computing (HPC) resources managed by the SLURM resource manager through an easy-to-use graphical user interface. Secondly, we enable asynchronous execution of workflows, a feature which guarantees that workflows will achieve reasonable resource utilization even when the scheduler cannot make use of a system’s full RAM and CPU resources. Thirdly, we propose a universal, platform agnostic JSON representation of workflows that allows platform-specific execution details to be abstracted away from the core scientific logic. Our work includes a custom executor plugin that supports translation of workflows from an external language, such as Nextflow, to our universal JSON format. Finally, we develop a graphical user interface to make our scheduler easy-to-use for non-technical users. When benchmarked on a bulk RNA sequencing workflow, these features reduced the cost and time requirements. We illustrated the merits of our cross-platform method using credit allocations from federally funded supercomputers.

## 1 Introduction

With the rapidly increasing size and scale of genomic, bioimage and spatial data, it is essential to develop scalable computational methods and infrastructure to process these increasingly large-sized datasets. Researchers have adopted a variety of techniques to cope with rapidly increasing scale of genomic data, mostly involving the use of cloud or local clusters for big biomedical data analyses. Supercomputers offer an alternative to these approaches as academic researchers can request credits and/or allocations from federally funded supercomputers and obtain the on-demand and high performance advantages of cloud computing without the onerous data egress costs for large datasets ($85 per TB transferred from AWS). Our key idea is to develop methods and tools to enable biomedical scientists to effectively leverage both commercial cloud and HPC for big data analysis. Effectively leveraging HPC and cloud computing requires significant technical knowledge. Users may wish to deploy the entirety of a bioinformatics workflow to a particular platform or may instead deploy different stages across different platforms so as to minimize cost or enhance parallelism.

Here, we present a scientific workflow scheduler based on the Temporal.io framework [1]. A major obstacle is that using supercomputers requires significant technical knowledge to adapt a bioinformatics workflow to execute on a supercomputer. We have implemented a graphical user interface (GUI) that allows users to take an existing workflow and specify which steps of the workflow should be executed on the supercomputer. This allows users to reduce execution time and optimize costs without changing the workflow specification or learning the technical details of supercomputer schedulers. Through a universal interchange JSON representation of workflows, we develop a custom executor plugin that allows users to specify workflows in a platform-agnostic format that focuses on the core logic and is compatible with external languages. We demonstrate the merits of our approach by benchmarking a bulk RNA sequencing workflow, showing appreciable makespan improvements (between 13% and 22%) when using asynchronous execution constructs. Additionally, we show how cross-platform scheduling of the bulk RNA sequencing workflow across local compute resources and remote HPC clusters can reduce consumption of compute credits, allowing better scaling of data-intensive bioinformatics pipelines. In one benchmarking result, we reduce the HPC credit consumption of a Bulk RNA sequencing pipeline by 25% simply by performing adapter trimming locally

### 1.1 Related Work

In scientific fields where large datasets are common, researchers commonly rely on Scientific Workflow Management Systems (SWMSs) to automate and standardize the processing of data. SWMSs such as Nextflow [2], Snakemake [3], Galaxy [4], Taverna [5], and the BioDepot Workflow Builder (BWB) [6] enable users to compose pipelines consisting of many independently-developed pieces of software. Nearly all SWMSs implicitly or explicitly represent workflows as directed acyclic graphs in which nodes represent individual computational steps and edges represent data dependencies between steps.

A key design choice among SWMSs is whether to implement workflow construction via a domain-specific language (DSL, as in Nextflow and Snakemake) or through a graphical user interface (as in BWB, Taverna, and Galaxy). This divide is particularly salient in the bioinformatics sphere, because biologists with limited programming experience often constitute a large portion of users. In our view, graphical-based SWMSs lag behind DSL-based SWMSs in their support for two key features: execution location transparency and fault tolerance. By *execution location transparency*, we mean that the user can assign workflow steps to different computational environments or resource managers (e.g. local execution, SLURM, etc.) without needing to encode file transfers or resource manager interfaces in the workflow definition. While Nextflow and Snakemake (as of version 8) support a variety of *executors*, including SLURM, AWS Batch, and Kubernetes, and automatically stage data dependencies on the relevant platforms [7] [8], support among graphical SWMS is much more limited. BWB provides no support for remote execution unless explicitly written into the workflow, and, while Taverna allows transparent submission of jobs to certain grid environments, it does not support modern common managers such as SLURM or AWS Batch nor does it allow for automatic file staging [9]. Galaxy provides limited support for cloud-based resources and HPC, though it has several limitations, most notably the requirement that resource managers be specially configured to interface with Galaxy using either Cloudman or a remote CLI, which is not suitable for certain types of remote HPC clusters [10]. By *fault tolerance*, we refer both to the SWMS’s ability to recover from transient faults such as network errors or job pre-emption and the ability to resume workflow execution from a valid state (checkpointing). Galaxy and Taverna each support transient fault tolerance through a backoff-retry model but offer no checkpointing support, while BWB has neither checkpointing nor a formal retry model. This contrasts once again with Nextflow and Snakemake, each of which allow resumption of workflows from particular checkpoints [9].

Temporal is an open-source workflow engine with an emphasis on fault tolerance and durable execution [1] [11]. Temporal workflows are defined as code in any language for which Temporal’s maintainers provide an SDK. Temporal provides substantial fault tolerance guarantees by tracking a workflow instance’s history on a central server; this history includes the workflow’s inputs, signals and updates received from outside the workflow, activities invoked by the workflow and their results, and child workflows initiated during workflow execution. Temporal enforces the key constraint that workflows are deterministic, meaning that they produce behave identically when presented with the same event history; non-deterministic behavior must be executed within workflow activities. This allows the temporal server to serialize and reconstruct the state of a running workflow at some later time by *replaying* its event history; in a *replayed* workflow, Temporal will not re-execute completed activities, running only the workflow code needed to process their results and restore state. This allows Temporal to periodically retry failed workflows with some backoff. Likewise, users can suspend workflow execution while awaiting signals or updates from outside the workflow, and the workflow state will be quickly restored based on the event history once the relevant signal occurs, without a worker needing to track workflow state in the interim. These features, which Temporal often describes as *durable execution*, allow for interactive and long-running workflows, in contrast to most SWMSs.

Temporal distributes workflow execution through a publish-subscribe model. A temporal server stores running and pending workflows and activities, each of which is associated with a particular queue. Temporal workers periodically poll the server for unscheduled activities and workflows and begin executing them. Furthermore, if a workflow’s execution is suspended for a significant amount of time (e.g. by waiting on a long-lasting signal), a worker may cease processing a given workflow, allowing it to be rescheduled on a different worker on the same queue once the workflow’s state changes. If no workers are listening to a particular queue at the time an activity is submitted to it, the workflow simply suspends execution until a worker has processed the activity. This model ensures that Temporal workflows are impervious to worker failures and relieves the server from needing to track available workers at any given time.

Despite its prevalence in industry, Temporal has not seen widespread adoption in bioinformatics, for two main reasons. The first is the lack of a SWMS-style DSL. With tools such as Nextflow or Snakemake, users only require basic knowledge of shell scripting to begin writing bioinformatics pipelines, and, even lacking this, they can select from a wide variety of open-source bioinformatics workflows in those DSLs (e.g. nf-core). By contrast, implementing workflows in temporal requires substantial knowledge of Temporal’s architecture and SDKs. Second, Temporal’s scheduling mechanisms are poorly-suited for computationally-intensive scientific workflows. Temporal provides only very coarse guarantees regarding scheduling. Its default behavior is to reserve a number of “slots” for each task queue to which a worker subscribes and to accept new tasks from the relevant queue until each is filled. Users may configure the maximum number of concurrent tasks and the number of slots per queue. Temporal offers no way to specify the expected memory or CPU usage of jobs within a particular task queue, nor does it allow users to specify task priorities. Furthermore, because Temporal reserves slots before polling even begins and because workers have no knowledge of a task queue’s contents before polling, it is impossible to reserve system resources only after a job is ready to execute. These present substantial limitations for scientific workflows, where the memory and CPU requirements of workflow steps can vary substantially [12]. Temporal does provide an alternative option in which users can specify target RAM and CPU requirements for a worker, allowing workers to limit the acceptance of new tasks while approaching the limit; however, this too is unsuitable for scientific workflows, where a single task can consume all of a worker’s available memory or CPUs.

### 1.2 Our Contributions

This work encompasses three main contributions relative to the existing literature in workflow design. The first is the introduction of workflow language constructs to enable asynchronous execution of scientific workflows. As shown in the work below, the ability to begin executing a successor step in the workflow as soon as its predecessor has processed a single sample, rather than waiting for all samples to process, and the ability to constrain the number of samples processed at a given time allows the user to introduce optimizations that would be impossible in popular Scientific Workflow Management Systems (SWMSs) such as Nextflow [2] or snakemake [3]. Secondly, our scheduler allows users to transparently distribute different workflow steps to different computational environments, including both to temporal workers running on users’ own servers of the cloud and remote SLURM clusters. The use of the Temporal framework and its guarantees of durable execution render workflows executed by the scheduler resilient to worker failure. Finally, the development of a universal interchange JSON format for workflow representation that allows the specification of workflows in a platform-agnostic manner.

We present a new graphical SWMS which offers both execution location transparency and greater degrees of fault tolerance compared to existing platforms. Our platform supports existing Nextflow and BWB workflows once provided with minimal annotations (resource requirements and jobs’ input and output files). Users may execute workflows on local hardware, but they may also scale their workflows through the addition of new compute nodes (workers) or the use of external resource managers (e.g. SLURM). Using a graphical user interface, users can easily map workflow steps to diverse computational environments, allowing considerable optimizations in workflow cost and execution time with minimal technical knowledge requirements; figure

**??** displays the interface. The backend to this SWMS is built atop the Temporal project [1], a workflow management system which tracks workflow history in a centralized server. By making use of Temporal’s default interfaces for backoff and retrying of workflow activities, we achieve transient fault tolerance guarantees similar to most other SWMSs, including the recovery from worker failure. Additionally, by retrieving the history of partially run workflows, we support the pausing and resumption of running workflows.

## 2 Materials and Methods

### 2.1 Workflow Interchange Format

The contemporary bioinformatics landscape is characterized by a diverse and powerful ecosystem of workflow languages and execution engines, including Nextflow, CWL, WDL, and Snakemake[13]. While each system offers distinct advantages, this technological heterogeneity has created a significant challenge: a fundamental lack of interoperability. Workflows defined in one language are not directly compatible with the execution engines of another, creating a fragmented ecosystem that acts as a considerable barrier to reproducible research. The manual translation of intricate logic, dependencies, and environment configurations between systems is not only labor-intensive but also introduces a high risk of subtle errors that can alter scientific outcomes, directly undermining the core tenets of reproducibility.

To address this challenge, we propose a universal, platform-agnostic solution centered on the representation of all workflows as directed acyclic graphs (DAGs). This graph-based model serves as a common intermediate representation, abstracting the core scientific logic of a workflow from the implementation-specific details of any particular language. In this model, a complex analysis is deconstructed into its fundamental components. A standard RNA-Seq pipeline, for example, would be represented as a graph where a quality control step using FastQC is one node, and a subsequent alignment step using the STAR aligner is another. A node encapsulates all critical information required for its execution: the specific software tool, its version, its parameters, and its computational resource requirements like CPU and memory. The data dependencies that dictate the order of operations are represented by edges. An edge connects the FastQC node to the STAR aligner node, signifying that the trimmed and quality-controlled FASTQ files produced by the former are the required input for the latter. This graph structure provides a canonical, language-independent blueprint of the entire scientific procedure. For the data serialization of this graph model, we selected JavaScript Object Notation (JSON) due to its ideal characteristics for data interchange. Its lightweight, human-readable syntax is easily parsed by virtually any programming language, making it a universal standard for communication between disparate systems. To ensure the structural integrity and correctness of this representation, we employ JSON Schema as a formal specification. The schema acts as a strict contract that defines the precise structure of the workflow graph. It enforces validation rules, such as ensuring that every node specifies a container or that memory requirements are defined in a consistent format. This prevents malformed or incomplete workflow definitions from ever being submitted for execution, thereby averting the significant waste of computational resources and researcher time.

This approach allows platform-specific execution details to be abstracted away from the core scientific logic. In collaborative, multi-institutional projects, it is common for participating sites to utilize heterogeneous computing environments, ranging from local Slurm clusters to various cloud platforms like AWS Batch.

A single, workflow definition can contain custom annotations, such as nextflow:label for a Slurm-based cluster or bwb:container for a cloud-based engine. These annotations provide specific execution hints to different target environments without altering the shared, core workflow structure, making the representation truly “execution-aware” and portable.

The primary mechanism for translating workflows from an external language, such as Nextflow, into our universal JSON graph representation is a custom executor plugin. This strategy leverages the native extensibility of the source workflow system to capture execution details dynamically, a more robust approach than attempting to statically parse a complex workflow script. The process is initiated when a bioinformatician launches their pipeline with a specific flag that activates the plugin for an “observational” run. As the source engine prepares to execute each task, the plugin’s custom hook intercepts lifecycle events. It captures the complete context for that task: the exact command-line script, the specified container image, unique identifiers, and the precise paths to all input and output files. This dynamic, on-the-fly construction of the graph ensures a true and complete representation of the pipeline’s execution logic. The resulting JSON graph serves as a complete, platform-agnostic blueprint that our scheduler can ingest and execute, effectively decoupling the workflow’s definition from its execution and enabling advanced capabilities like fault tolerance and asynchronous processing.

**Fig 1:**
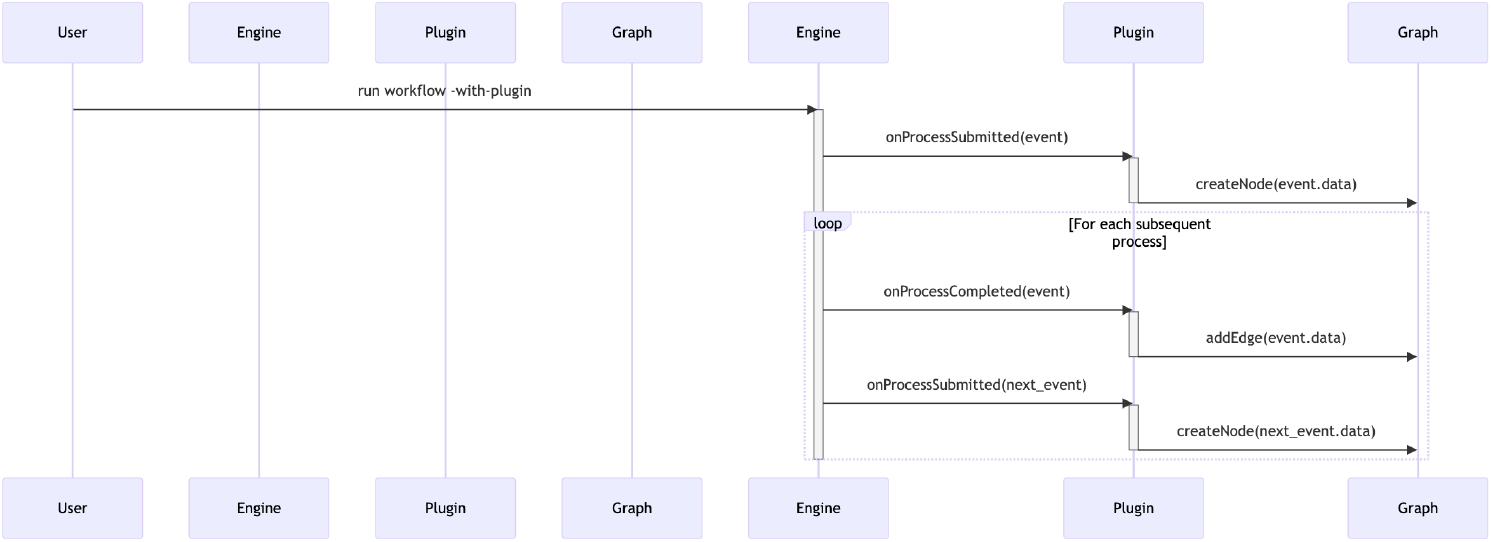
The diagram illustrates the interactions over time between lifelines representing the User, Nextflow Engine, Custom Executor Plugin, and JSON Graph. Messages depict the sequence: the User initiates an observational run, the Engine emits lifecycle events (onProcessSubmitted, onProcessCompleted), which are intercepted by the Plugin to progressively build the JSON Graph object.

### 2.2 Workflow Definitions and Configuration

The scheduler interprets workflow definitions based on a JSON format closely modeled after BWB’s workflow format. This similarity allows us to easily adapt existing BWB workflows into the scheduler’s format, but the format is sufficiently general to express workflows from other SWMSs. We hope to eventually create a workflow “interchange format” based on a superset of the existing format in order to support Nextflow and Snakemake workflows as well.

A workflow definition specifies an abstract DAG in terms of nodes and links. Each node corresponds to a unique identifier, a docker image, estimated CPU and memory requirements, and a set of default container, while each link transfers an output parameter of one node to serve as the input to another. Once the source node of a given link completes execution, the value of the link’s source node parameter generated by that run of the source node will be transferred over the link and will overwrite the default value of the sink node’s parameter. The value sent from the source node may be hardcoded within its parameters or may instead be dynamically set at runtime.

Workflow definitions can also specify iterable and asynchronous behavior within workflow nodes. An *iterable node* is one which executes multiple commands given a single set of inputs. A common use case involves executing a command once for every sample in a dataset, though the format inherited from OWS also supports more advanced use cases, such as iterating over every pair of associated paired-end FASTQ files based on a particular naming convention. Building on this, our workflow format introduces the concepts of *asynchronous* and *barrier* nodes. *Asynchronous nodes* are a subset of iterable nodes which generate a new set of outputs for each command executed. This allows them to trigger successor nodes once for every iteration of the source node such that successors are not blocked while waiting for their predecessors to process every sample in a dataset. Users may additionally identify a descendant of an asynchronous node as its “barrier”; this means that the barrier node will wait until all ancestors of itself which are descendants of the asynchronous node have completed an iteration corresponding to each iteration of the asynchronous node before the barrier node itself will generate a command and begin execution. We call these intermediate nodes which are descendants of the asynchronous node and ancestors of the barrier the *asynchronous descendants* of the asynchronous node. The workflow definition allows users to place an upper limit on the number of concurrent executions of an asynchronous node whose asynchronous descendants have not yet completed, which we call the *maximum asynchronous concurrency*; once this limit is met, the scheduler will exclusively focus on scheduling asynchronous descendants of executed asynchronous nodes rather than scheduling new asynchronous nodes. This is useful for controlling allotments of disk space which are held and released across multiple nodes within an asynchronous block.

### 2.3 Scheduler Implementation

Execution of a workflow involves five components, three essential and two optional: a client, which submits a workflow request; the Temporal server, which receives this workflow request and assigns it to a worker; a set of temporal workers, who receive and process the workflow and its constituent activities; an optional S3 server (supporting any implementation of the S3 API, including open-source options such as minio) to coordinate file sharing between workers and store final results; and optional external executors such as SLURM, which may receive and execute particular workflow tasks in lieu of the temporal workers. Figure 2 provides a diagram of these components.

**Fig 2:**
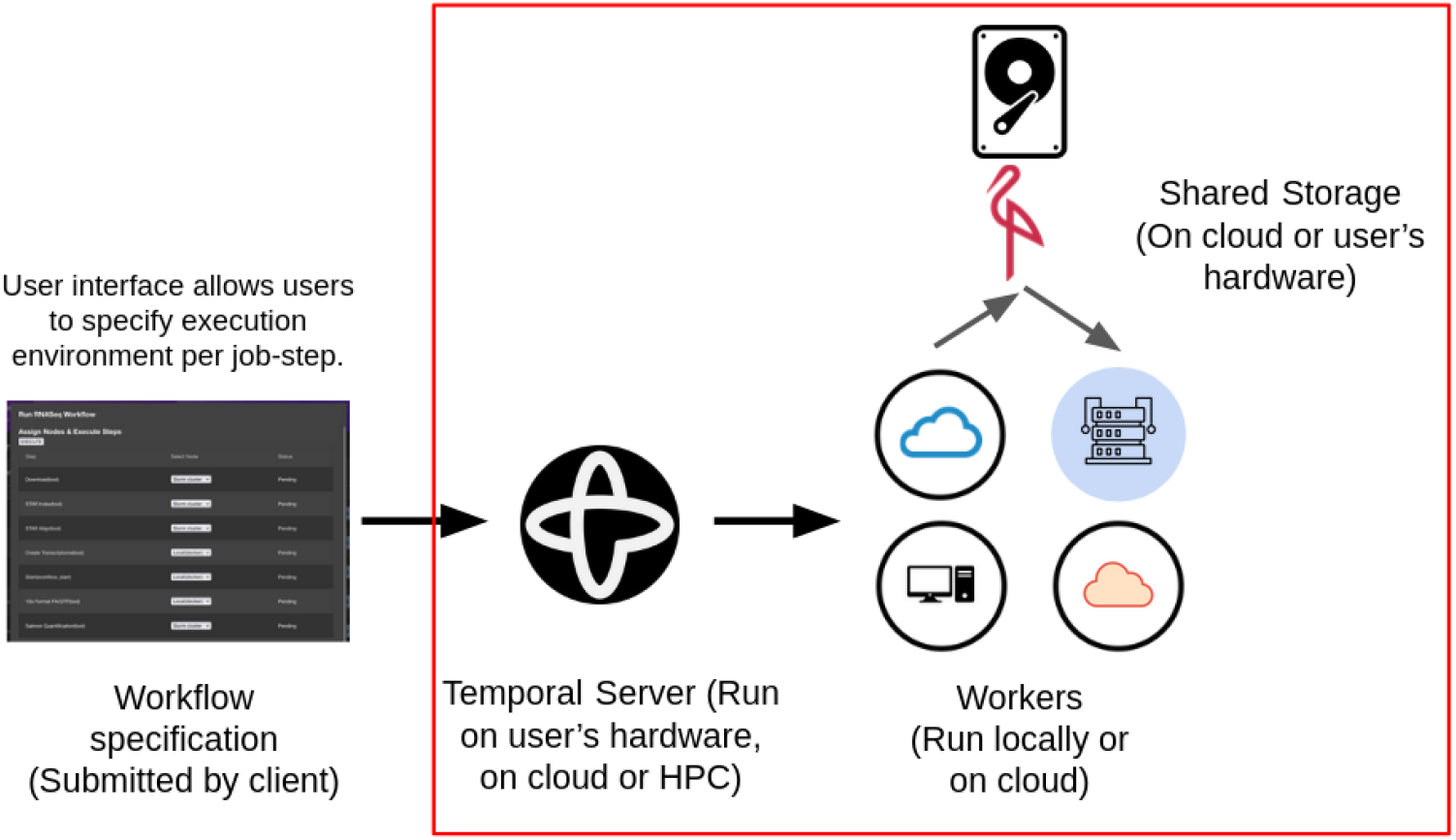
Architecture diagram of the scheduler. Those items in the red box were designed as part of this thesis. The client and user interface remain works in progress and are outside the scope of my work.

**Fig 3:**
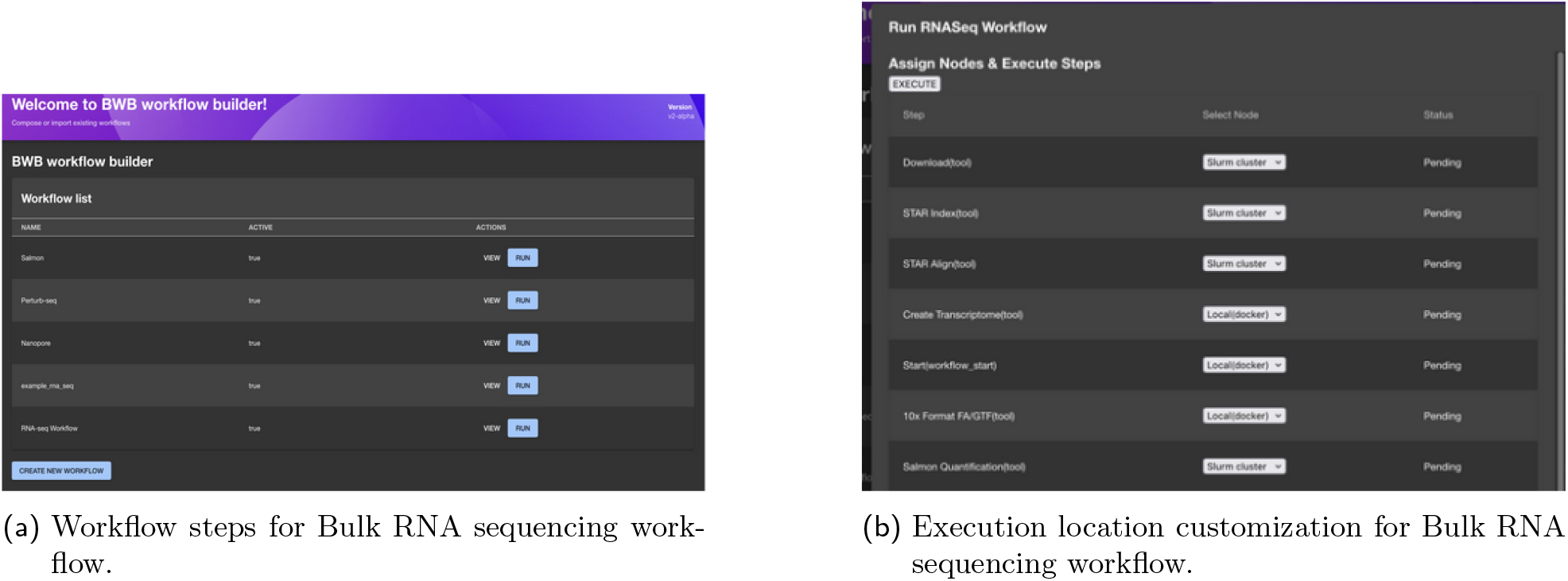
User interface for our SWMS.

**Fig 4:**
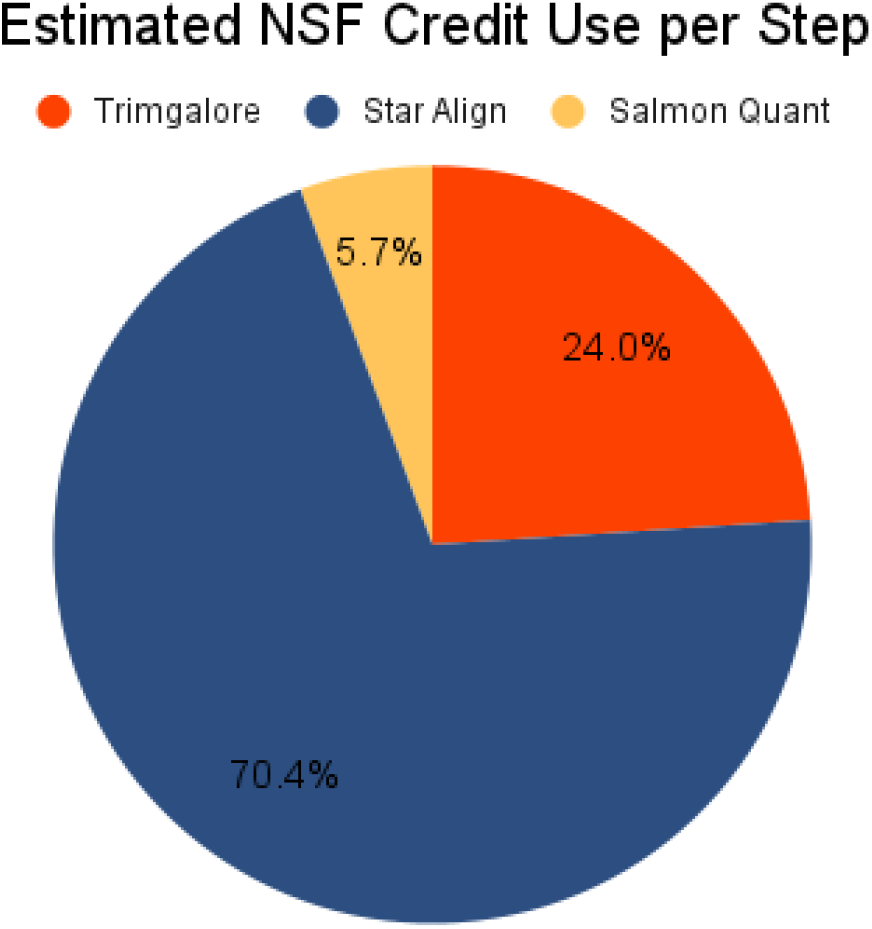
Estimated credit use per step of pipeline based on conversion of SLURM job runtime to CPU hours.

Our implementation distinguishes between two types of temporal workers – *scheduler workers* and *regular workers*. A *regular worker* executes the actual commands of a workflow. It has a unique queue name and is initiated by the user with limits on RAM, CPU, and GPU usage that will be respected during task scheduling. When the scheduler assigns a command to a regular worker, activities in order to stage the command’s prerequisite files, execute the command itself, and upload the results are routed to the worker’s temporal queue. In contrast, *scheduling workers* execute tasks that do not require resource grants, such as the generation of workflow commands, the tracking of workflow inputs and outputs, and the assignment of workflow jobs to workers. As none of the tasks assigned to *scheduling workers* depend on file locality, they share a common queue. This division ensures that tasks responsible for a running workflow’s control flow do not wait in Temporal queues behind workflow jobs.

As discussed in the literature review, Temporal’s default publish-subscribe model, which relies on slots-based rate limiting, is a poor fit for cases involving variable resource requirements between jobs. We implement a push-based job system as part of the core workflow logic while still obeying Temporal’s requirement for deterministic workflow behavior. In our system, the main workflow initiates a child workflow responsible for the assignment of outstanding jobs to workers. This child workflow is submitted to the shared queue for scheduler workers. Each job, specified by a unique ID alongside a required allotment of memory, CPU cores, and GPUs, is communicated from the main workflow to the child workflow via a temporal signal, and the child workflow’s responses are communicated back to its parents via signals. Likewise, regular workers announce their existence to the child workflow via temporal signals. Upon receiving a signal from a worker, including its unique queue ID, the scheduling child workflow starts a heartbeat activity on the subscribed worker’s unique queue. (More specifically, each regular worker maintains a distinct queue for heartbeat activities so as to ensure that these activities are not preempted by workflow commands in the temporal slot system.) Using a temporal primitive, this heartbeat activity will continue indefinitely so long as the worker communicates its liveliness to the server every 20 seconds; failing that, a workflow exception will be triggered, and the child workflow will register the worker as dead. It will then attempt to reschedule the heartbeat activity to the worker queue using an exponential backoff. Periodically, the child scheduling workflow will execute an activity to assign outstanding job requests to workers, tracking the resources remaining on each worker.

This periodic logic motivates our decision to implement the scheduling workflow as a child workflow rather than an integral part of the main temporal workflow; temporal workflows have an upper limit on the size of their event histories, which requires them to periodically “Continue-As-New”, an operation which involves terminating ongoing processes and reconstructing workflow state in a new workflow. By segregating periodic logic into child workflows, we limit the size of the main workflow’s event history and circumvent the need to stop and resume ongoing workflow commands once the job history grows unmanageably large. When the scheduler child workflow’s event history reaches some pre-defined size limit (by default, one-fifth of the maximum value), it cancels its ongoing heartbeat activities, serializes its state, and invokes the “ContinueAs-New” operation, rescheduling all heartbeat activities within this new workflow. Because the execution of the child scheduling workflow is determined entirely by temporal signals (from both workers and its parent workflow) and the results of activities (including exceptions triggered by failed heartbeat activities), it behaves deterministically and can be replayed by the temporal server in case the parent workflow fails and is resumed.

We treat scheduling as a vector-packing problem in which each worker has a fixed allotment of CPU cores, GPUs, and memory and where each job will occupy some subset of those resources. Our approach to scheduling consists of a simple first-fit decreasing heuristic: we loop through all available workers, then loop through all outstanding job requests ordered by their most demanding resource request as a fraction of the corresponding bin dimension, and assign each job request to the first worker capable of accommodating it. The aim of such an algorithm is to assign all available jobs to as few workers as possible, thereby maximizing resource utilization and preventing resource fragmentation. Garey et al. (1976) have shown that this algorithm achieves a competitive ratio of of 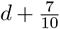 relative to the offline version, where *d* is the number of dimensions, which makes it suitable for low-dimensional cases [14]. In practice, most workflow jobs will operate with *d* = 2, since only a handful of jobs require GPU grants (in which case *d* = 3).

The choice of an online algorithm is motivated by several factors. Firstly, our workflow format provides support for dynamic workflow definitions, in which the precise set and number of commands to be executed is not known at the start of workflow execution and may be updated during the workflow runtime. (This enables workflows to poll for and process arbitrarily large numbers of files as they appear.) This approach is fundamentally at odds with workflow scheduling algorithms such as HEFT, which demand comprehensive knowledge of the workflow graph before execution. Secondly, we may wish to eventually support an architecture in which workers are used by multiple users concurrently; with an online scheduling algorithm already in place, this could be accommodated almost instantly by switching the scheduling child workflow into a fully fledged Temporal workflow which may be signaled by several running workflows simultaneously rather than accepting requests exclusively from its parent workflow. Finally, heuristic-based bin-packing is a substantially simpler algorithm to implement than most offline scheduling algorithms.

One of the major aims of our scheduler is to allow users to easily distribute workflows across multiple execution environments, executing jobs both on their own hardware, on the cloud, and on HPC. Execution on the cloud requires no special treatment; users can provision on-demand compute instances from any major cloud provider and run a temporal worker, using the same temporal server to which the client submits workflows. By contrast, execution on multi-user HPC clusters typically requires interfacing with a distinct resource manager such as SLURM or Kubernetes. Furthermore, users on these systems often lack the relevant permissions to install the dependencies necessary to run a temporal worker; the only permissions we can generally assume are the ability to submit jobs, either through a remote shell on a login node, as in most SLURM systems, or through some API, as in Kubernetes, HTCondor, etc. To help overcome differences in resource managers’ interfaces, we introduce the abstraction of an “executor”. An executor is an abstract interface which provides general functionalities to run workflow jobs, retrieve their results, transfer file dependencies, and handle resource allocations on a particular resource manager. At present, we have only implemented two executors: one for execution on local temporal workers and another for executing jobs on remote SLURM clusters. However, the interface is sufficiently general that it may be easily extended to other resource managers in the future.

As with the scheduling of jobs to activities, the submission of workflow jobs to remote resource manager relies on a child workflows. Our SLURM executor submits workflow jobs to a remote SLURM cluster via an SSH connection. The management of this SSH connection introduces some difficulties. We cannot interact with networked communication within the Temporal workflow, as this would violate determinism constraints. Likewise, we cannot open a new SSH connection within each activity, since this could cause the SLURM login node to refuse connections due to an upper bound on concurrent SSH sessions for a single user. Our solution is to delegate the management of the connection lifetime to a Temporal worker. We create a special class of worker which maintains an SSH connection with a particular SLURM cluster and whose queue name is uniquely determined by the username and endpoint associated with this SSH session; the SLURM executor then assigns temporal activities to this queue, so that activities can reuse the existing SSH connection already open on the worker. Failure of the SSH connection results in temporary failure of the worker, causing the SLURM child workflow to block until a queue corresponding to the relevant cluster becomes available. It is incumbent upon the client to ensure that a worker with a connection to the given SLURM cluster is available when a workflow requests it. To make this simple, we provide a script to users which takes a workflow definition, submits it to the temporal server, automatically establishes all workers necessary to complete said workflow’s execution, and terminates these workers once the workflow has completed execution. We anticipate that our user interface will provide similar functionality.

Our executor for SLURM clusters functions primarily by sending and receiving signals to and from a child workflow. The main workflow assigns jobs to SLURM through a signal to the child workflow, and the child workflow returns results and logs through a signal to its parent. Requests to execute a command on the SLURM cluster include the job’s estimated resource requirements and SLURM-specific configuration information (e.g. desired partition). Upon receiving a request from the main workflow, the child workflow submits an activity to the queue of the worker responsible for maintaining the SLURM connection, which in turn submits the command to the SLURM cluster via an sbatch script. The child workflow also periodically submits a polling job to this queue, specifying all outstanding SLURM jobs. The worker responsible for maintaining the connection with said cluster then submits an sbatch command which retrieves the state of all jobs in the workflow’s activity parameters, as well as the standard output and standard error of completed activities. It is this polling activity which necessitates the segregation of SLURM handling into a child workflow, and said child workflow periodically invokes the continue-as-new API to prevent event history overflows.

Because Docker adopts a lax approach towards user privileges, multi-user SLURM systems generally require users to use Singularity instead. Our software provides automatic conversion from docker to singularity via the Open Countainer Initiative (OCI) format.

SLURM clusters may enforce particular constraints on jobs’ resource requests. For instance, a cluster may require that jobs’ request for memory not exceed a certain fraction of their requests for CPUs. Likewise, SLURM clusters will terminate jobs which exceed their memory allotments. To cope with these complexities, we allow users to override the memory, CPU, and GPU requests specified in their workflow definition within their workflow configurations. Should they fail to do so, we submit jobs to SLURM with CPU and GPU requests unmodified and with memory requests augmented by 20%, in order to provide a cushion for jobs which slightly surpass RAM estimates. (Snakemake does the same.)

### 2.4 Graphical User Interface

While a standardized interchange format is a powerful solution for machine-level interoperability, the direct manipulation of workflow definition files presents a steep learning curve for many domain experts who are not computational specialists. The potential for subtle errors in parameter specification or file path syntax can lead to failed analyses, wasting significant computational resources and researcher time. This accessibility barrier has been a primary motivator for the development of user-friendly graphical interfaces in widely used bioinformatics platforms like Galaxy, which are designed to make complex computational tools accessible to all researchers, regardless of their programming expertise. Our goal is to provide a similar level of accessibility by abstracting away the underlying syntax of the workflow definition through an intuitive visual interface.

## Results

### 3.1 Use case: Bulk RNA sequencing data

To demonstrate the benefits of execution location transparency, we conduct some simple benchmarks on the MORPHIC Bulk RNA sequencing workflow, a standard pipeline for alignment of Bulk RNA reads against a reference genome modeled closely after the nf-core equivalent. The pipeline includes a number of preliminary and preprocessing steps, such as the download of genome and transcriptome annotations, the download of input FASTQ files from an S3 bucket, and the building of a genome index, all of which may be elided if the relevant files are already present on the system. The bulk of the pipeline’s computational time is spent in three steps: adapter trimming using Trimgalore (a wrapper around CutAdapt [15]), alignment to the genome and transcriptome using STAR [16], and quantification using Salmon [17]. Each of these steps are iterable, meaning that they execute once for every sample (meaning once for every single-ended FASTQ file or once for every pair of paired-end FASTQ files) in the dataset. At the end of this process, a lightweight bash script constructs a counts table that quantifies gene expression from the salmon results. The alignment step is by far the most intensive, requiring roughly 32 GB of memory in order to store a genome index, as well as 16 threads. By contrast, trimming requires only around 4GB of memory and 4 threads, and quantification requires only 8 GB with 4 threads.

We benchmark the Bulk RNA sequencing pipeline on the NSF Bridges 2 supercomputer. We obtained an NSF grant of 200,000 credits on this supercomputer. Credit consumption is billed based on CPU usage, with one CPU hour corresponding to a single credit in most partitions of the SLURM cluster. (Pricing may vary in high-demand clusters, particularly those with GPUs.) Users may also expend credits to reserve space on the cluster’s highly parallel Lustre filesystem, with one credit corresponding to a single gigabyte of storage. We obtained a 10 terabyte allotment of disk space. For all benchmarks, we submitted jobs to the RM-shared partition, a multi-user partition consisting of 396 nodes, each with 128 CPU cores and 256 GB of RAM. This partition requires that user job requests have a memory-to-CPU ratio not exceeding two gigabytes. In cases where the RAM and CPU requirements used for local execution did not obey this constraint, we simply increased CPU requirements to the minimum value which would achieve such a ratio.

We tested three configurations of the pipeline: the first executed all steps on an AWS d3en.8xlarge instance; another executed all steps on the NSF Bridges2 supercomputer (full HPC); the final configuration executed the trimming step locally while executing all other steps on the supercomputer (hybrid HPC). Performing trimming locally is a desireable optimization, because the trimming step is disk-bound with low RAM and CPU requirements; on multi-user filesystems such as Lustre, this can result in substantial latency and CPU underutilization, costing credits for a step that could be easily executed on a local server.

Two datasets from the MORPHIC program were used for benchmarking. The first aims to study the effects of knocking out two transcriptor factors in brain and other support cells (publicly available from NCBI GEO with accession number GSE287843 [18]). This dataset consists of 28 sets of Illumina-sequenced, paired-end FASTQ files which, when compressed, occupy 181 GB of disk space. The second dataset consists of 90 samples that aim to phenotype iPSC-derived trophoblast lines that contain homozygous null alleles for 7 transcription factors expressed within the extra-embryonic lineage (GEO accession GSE288288 [19]). These benchmarks use the published GSE287843 [18] and GSE288288 datasets [19] that aim to study the effects of knocking out two transcriptor factors in brain and other support cells.

Table 1 gives raw results in terms of task runtimes, while table 2 summarizes relative runtime and cost savings. By executing the trimming step locally, we realize a reduction in HPC credit consumption of roughly a quarter in the hybrid pipeline. Additionally, running the pipeline on HPC achieves a speedup of between 6 and 9 times relative to AWS with the full HPC pipeline configuration (between 3.8 and 4.3 times with the hybrid HPC configuration).

**Table 1:** Comparison of runtime and across AWS, full HPC, and hybrid HPC for two datasets.

| Dataset | AWS Runtime | Full HPC Runtime | Hybrid HPC Runtime | Full HPC Credits | Hybrid HPC Credits |
| --- | --- | --- | --- | --- | --- |
| GSE287843 | 11h 35m | 01h 53m | 03h 01m | 352 | 261 |
| GSE288288 | 25h 36m | 02h 44m | 05h 55m | 872 | 671 |

**Table 2:** Benchmarking results across two bulk RNA-seq datasets. AWS compute cost is calculated by multiplying the execution time of the entire workflow by the hourly rate of an AWS EC2 d3en.8xlarge instance at $3.996 per hour. Speedup is calculated by the ratio of the execution time using AWS EC2 d3en.8xlarge instance over that of full HPC mode using Bridges-2. Credit savings is calculated by the HPC credits difference between the hybrid and full HPC mode over credits used by full HPC.

| Dataset | Dataset Size | AWS Cost (\$3.996/hr) | Speedup (AWS vs Full HPC) | Speedup (AWS vs Hybrid HPC) | Credit Savings |
| --- | --- | --- | --- | --- | --- |
| GSE287843 | 181 GB | \$46.31 | 6.13x | 3.83 | 25.85% |
| GSE288288 | 485 GB | \$102.30 | 9.37x | 4.32 | 23.05% |

By providing users with a simple, UI-based approach to execution location customization, we anticipate that they can achieve similar savings in credits when executing pipelines on the cloud or on HPC resources. Additionally, because of our integration with the Temporal project, our software can handle the numerous faults inherent to distributed execution of workflows.

## Conclusions

We anticipate that the research presented in this paper will reduce the technical knowledge required for the scaling and optimization of HPC-based bioinformatics pipelines. The introduction of a graphical SWMS with the ability to customize execution location of individual job steps and transparent file transfers allows users to map workflow tasks to appropriate environments with just a few clicks. Likewise, the integration of our universal interchange format with the UI will enable standardized pipelines for customization and deployment of platform-agnostic workflows.

These advantages are particularly relevant in light of the growing sizes of multi-omics data generated for biomedical research. The analyses of such data are often complex and require substantial computing resources. The use of HPC and federally funded supercomputers, particularly when using optimized analytical pipelines based on cross-platform execution, presents a cost-effective alternative for academic researchers to scaling these pipelines on commercial cloud. Substantially, the NSF Bridges 2 supercomputer used for benchmarking in this paper does not charge based on data ingress or egress, eliminating a major cost of cloud execution. Assuming linear scaling of credit consumption relative to dataset size, our 200,000 credit allotment from the NSF would allow execution of 568 bulk RNA-seq datasets of a similar size to GSE287843 when executing all steps on HPC resources. When introducing cross-platform execution optimizations such as local trimming, this number grows to 766. In short, the use of publicly-funded supercomputers will help cope with the rising size of multi-omics datasets while minimizing cost to individual researchers.

## Data Availability

Bulk RNA-seq datasets used in this paper are publicly available from the NCBI Gene Expression Omnibus (GEO) repository with accession numbers GSE288288 [19] and GSE287843 [18].

## Acknowledgement

We would like to thank Professor Paul Robson and Professor Bill Skarnes at the Jackson Laboratory and NIH grant UM1HG012651 used for generating the MorPhiC bulk RNA-seq datasets. We would like to thank the data ingestion team at the European Bioinformatics Institute (Anu Shivalikanjli, Galabina Yordanova, Alexandros Orges Koci, Sandeep Selvakumar) for data collection, metadata annotation and brokering of the MorPhiC RNA-seq datasets.

We would like to thank Dr. Phil Blood at the Pittsburgh Supercomputing Center (PSC) for discussions. This work used Bridges-2 at PSC through allocation BIO230124: Exploring conducting bioinformatics analysis on HPC for the NIH Morphic project from the Advanced Cyberinfrastructure Coordination Ecosystem: Services Support (ACCESS) program, which is supported by National Science Foundation grants 2138259, 2138286, 2138307, 2137603, and 2138296.

## Authors Information

L.H.H. and K.Y.Y. are investigators in the Data Resource Administrative Coordinating Center (DRACC) of the NIH funded MorPhiC program [20].

